# RASA1-driven cellular export of collagen IV is required for the development of lymphovenous and venous valves in mice

**DOI:** 10.1101/2020.02.17.953364

**Authors:** Di Chen, Xin Geng, Philip E. Lapinski, Michael J. Davis, R. Sathish Srinivasan, Philip D. King

**Affiliations:** Department of Microbiology and Immunology, University of Michigan Medical School, Ann Arbor, MI, USA; Cardiovascular Biology Research Program, Oklahoma Medical Research Foundation, Oklahoma City, OK, USA; Department of Medical Pharmacology and Physiology, University of Missouri, Columbia, MO, USA.

## Abstract

RASA1, a negative regulator of the Ras-mitogen-activated protein kinase (MAPK) signaling pathway, is essential for the development and maintenance of lymphatic vessel (LV) valves. However, whether RASA1 is required for the development and maintenance of lymphovenous valves (LVV) and venous valves (VV) is unknown. In this study we show that induced endothelial cell (EC)-specific disruption of *Rasa1* in mid-gestation mouse embryos did not affect initial specification of LVV or central VV but did affect their continued development. Similarly, switch to expression of a catalytically inactive form of RASA1 resulted in impaired LVV and VV development. Blocked development of LVV in RASA1-deficient embryos was associated with accumulation of the basement membrane protein, collagen IV, in LVV-forming EC and could be partially or completely rescued by MAPK inhibitors and drugs that promote collagen IV folding. Disruption of *Rasa1* in adult mice resulted in venous hypertension and impaired VV function that was associated with loss of EC from VV leaflets. In conclusion, RASA1 functions as a negative regulator of Ras signaling in EC that is necessary for EC export of collagen IV, thus permitting the development of LVV and the development and maintenance of VV.

## Introduction

The Ras signaling pathway is a ubiquitous intracellular signaling pathway that is triggered by different growth factor receptors (GFR) in numerous cell types.^1, 2^ Ras is a small GTP-binding protein, tethered to the inner leaflet of the cell membrane, which switches between inactive GDP-bound and active GTP-bound states. GFR activate Ras through recruitment to membranes of one or more Ras guanine nucleotide exchange factors (RasGEFs) that eject GDP from the Ras guanine nucleotide-binding pocket, thus permitting Ras to bind GTP.^3^ Activated GTP-bound Ras induces different downstream signaling cascades including the mitogen-activated protein kinase (MAPK) and phosphatidylinositol 3-kinase (PI3K) signaling cascades that couple GFR and Ras to cellular responses.^4, 5^ Inactivation of Ras is mediated by Ras GTPase-activating proteins (RasGAPs) that increase the ability of Ras to hydrolyze bound GTP to GDP by several orders of magnitude.^6^

There are ten different members of the RasGAP family that each comprise of a catalytically active GAP domain together with one or more modular binding domains.^6^ Most members of the RasGAP family are expressed broadly in different tissues and cell types. Nonetheless, one RasGAP family member, p120 RasGAP (also known as RASA1), has emerged as a critical regulator of blood vessel (BV) and lymphatic vessel (LV) systems in both humans and mice. In humans, germline inactivating mutations of the *RASA1* gene cause the autosomal dominant blood vascular disorder, capillary malformation-arteriovenous malformation (CM-AVM).^7–9^ Recent evidence indicates that the development of BV lesions in CM-AVM is dependent upon the acquisition of somatic inactivating second hit mutations in the intact *RASA1* allele in endothelial cells (EC) or their progenitors during development. ^10, 11^ This is consistent with the variable phenotype and location of BV lesions in CM-AVM patients. In addition, LV abnormalities, including lymphedema, chylous ascites, chylothorax and LV hyperplasia have also been observed in some CM-AVM patients.^8, 9, 11–14^

Non-conditional disruption of the *Rasa1* gene in mice results in mid-gestation lethality (E10.5) as a consequence of abnormal vasculogenesis and developmental angiogenesis in which primitive vascular plexuses are remodeled into hierarchical arterial-capillary-venous networks.^15, 16^ In addition, disruption of *Rasa1* in neonatal and adult mice results in impaired retinal angiogenesis and pathological angiogenesis respectively.^17–19^ We recently demonstrated that in the absence of RASA1, endothelial cells (EC) are unable to export the extracellular matrix protein, collagen IV, which is major constituent of vascular basement membranes (BM). This inability to export collagen IV, accounts for the impaired developmental, neonatal and pathological angiogenic responses in the absence of RASA1.

Concerning the LV system, induced disruption of *Rasa1* in adult mice results in LV leakage in the form of chylous ascites and chylothorax.^18^ This can be explained on the basis that RASA1 is required for the maintenance of intra-luminal valves in collecting LV that normally prevent backflow of lymph fluid.^20^ Upon *Rasa1* disruption, lymphatic EC (LEC) are lost from LV valve leaflets at the rate of approximately one LEC per leaflet per week until a threshold point is reached where the valve is unable to prevent fluid backflow and is unable to function.^20^ RASA1 is also necessary for the development of LV valves in late gestation. Disruption of *Rasa1* just prior to LV valvulogenesis at E15.5 does not impact upon initial LV valve specification characterized by increased expression of the PROX1 transcription factor in valve-forming LEC.^20, 21^ However, all PROX1^hi^ valve-forming LEC undergo apoptosis shortly thereafter and valve development ceases.^20^

Lymph fluid is returned to the blood circulation via LV that connect with the blood vasculature at the junction of the internal jugular vein (IJV), external jugular vein (EJV) and subclavian veins (SCV) with the superior vena cava (SVC).^22, 23^ Lymphovenous valves (LVV) guard the four points of connection of these LV with the blood vasculature (two points of connection on either side of the animal) and, thereby, prevent backflow of blood into LV.^22, 23^ LVV first form at E12 at the point of contact of lymphatic sacs (LS) with the blood vasculature. LVV specification is marked by the up-regulation of PROX1 by both LEC on the lymphatic side and LVV-forming EC on the venous side of the nascent valve structure.^22, 23^ LVV development then proceeds through the successive stages of delamination, aggregation and maturation as LVV leaflets form and extend into the vein lumen; a process that is considered complete by E16.5.^22^ Mature LVV leaflets comprise of two different types of EC attached to a central extracellular matrix (ECM) core.^22^ EC on the lymphatic and venous sides of leaflets continue to express high levels of PROX1. LEC on the lymphatic side of leaflets express high levels of vascular endothelial growth factor receptor 3 (VEGFR3), the receptor for VEGF-C. In contrast, EC on the venous side of LVV leaflets express low levels of VEGFR3 while the rest of the venous endothelial cells are VEGFR3 negative. Since RASA1 is necessary for the development of LV valves, it is important to determine if it performs a similar function in the development of LVV and, if so, at which stage.

VV prevent backflow of blood and are required for unidirectional flow of blood in veins^24^. Dysfunction of VV has serious consequences that include chronic venous insufficiency, venous hypertension and deep vein thrombosis.^25, 26^ Development of VV in central veins, such as the IJV, EJV and SCV, is initiated at E14.5 and is completed by E16.5.^22, 27^ In contrast, VV in peripheral veins, such as the saphenous vein, develop postnatally.^27, 28^ VV specification, like LV valve and LVV specification, is marked by increased expression of PROX1 in valve-forming EC.^22, 24^ Mature VV are structurally similar to LV valves in that leaflets of both type of valve comprise of a single type of EC attached to a central ECM core. VV and LV valve leaflet EC both continue to express high levels of PROX1 and share several other molecular characteristics including high level expression of the FOXC2 transcription factor and alpha-9 integrin.^22, 24^ Given the similarity between VV and LV valves, investigation as to whether or not RASA1 is necessary for the development and maintenance of VV is also important.

In the current studies, we used different mouse models of RASA1 deficiency to demonstrate a role for RASA1 in the development and maintenance of LVV and VV. Furthermore, we established that RASA1 is necessary for the export of collagen IV from valve-forming EC, which explains its role in valve development. Studies are of potential relevance to an understanding of the pathogenesis of CM-AVM and are of additional significance to other cardiovascular diseases.

## Results

### RASA1 is not required for the specification of LVV and central VV

Specification of LVV occurs at E12 and is marked by increased expression of PROX1 in LEC and LVV- forming EC at the points that LS make contact with BV in the region of the junction of the IJV, EJV, SCV and SVC.^22, 23^ Therefore, to examine if RASA1 plays a role in LVV specification, we administered TM to pregnant *Rasa1^fl/fl^* mice carrying *Rasa1^fl/fl^* and *Rasa1^fl/fl^ Cdh5^ert2cre^* embryos at E10.5 and analyzed embryos at E12.5. The *Cdh5* promoter drives expression of the *ert2cre* transgene within EC specifically.^29^ As assessed by staining of frontal tissue sections for PROX1, VEGFR3 and CD31, induced disruption of *Rasa1* at E10.5 in *Rasa1^fl/fl^ Cdh5^ert2cre^* embryos did not affect LVV specification (Figure 1 A).

**Figure 1.**
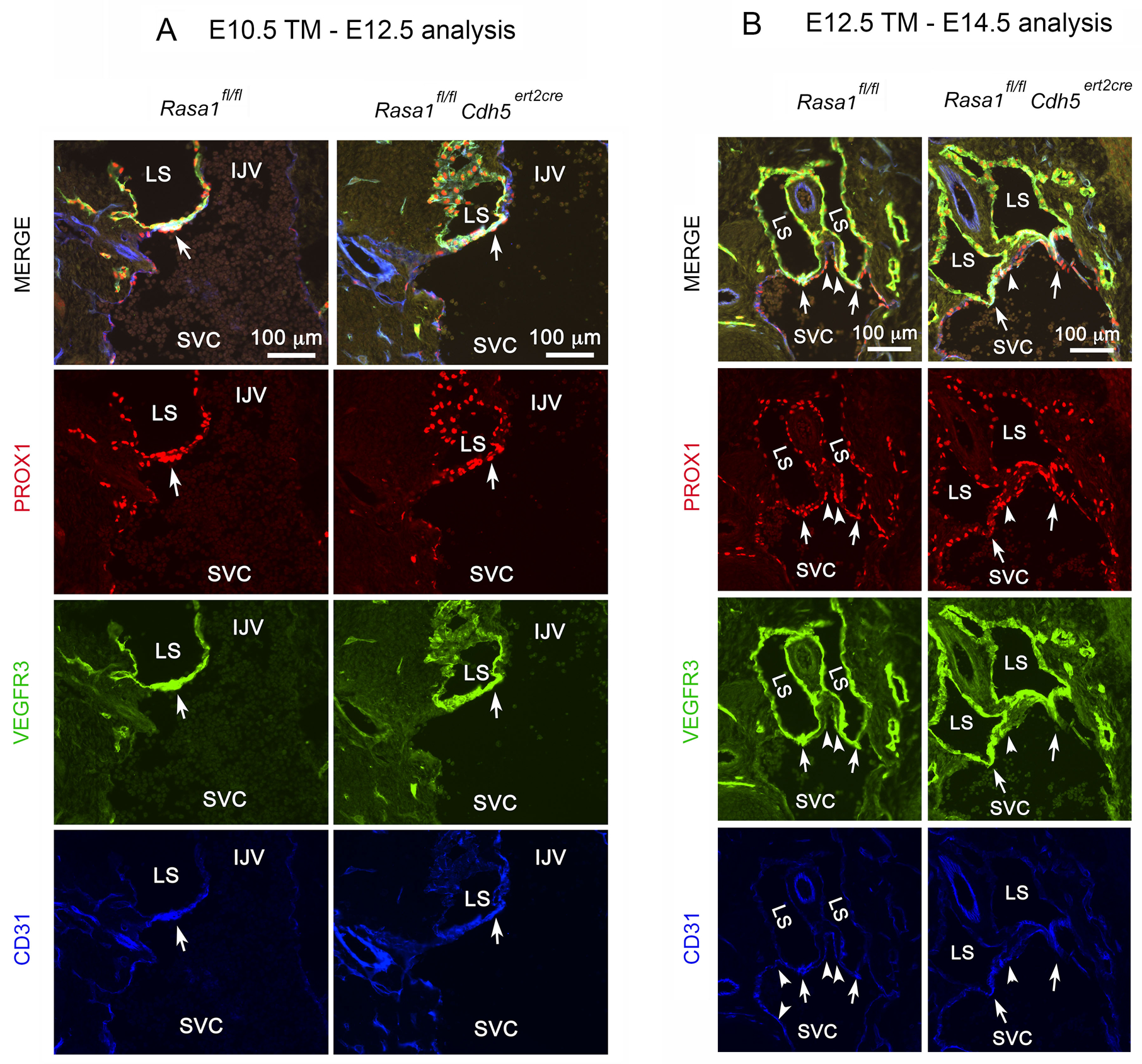
RASA1 is not required for the specification of LVV and central VV. **A** and **B**, *Rasa1^fl/fl^* and *Rasa1^fl/fl^ Cdh5^ert2cre^* embryos were administered TM at E10.5 or E12.5 and harvested at E12.5 or E14.5 respectively. Longitudinal sections through the neck region were stained with the indicated antibodies. LS, lymphatic sac; IJV, internal jugular vein; SVC, superior vena cava. LVV are indicated by arrows and VV by arrowheads. Note normal specification of LVV at E12.5 and VV at E14.5 (n=2 and 5 *Rasa1^fl/fl^* and *Rasa1^fl/fl^ Cdh5^ert2cre^* embryos respectively at E12.5 and n=4 and 5 *Rasa1^fl/fl^* and *Rasa1^fl/fl^ Cdh5^ert2cre^* embryos respectively at E14.5.

We also examined if RASA1 was required for the specification of central VV. Specification of the VV that guard the points of entry of the IJV, EJV and SCV into the SVC occurs at E14.5.^22, 27^ Similar to LVV and LV valve specification, VV specification is characterized by increased expression of PROX1 and VEGFR3 in VV-forming EC. Therefore, to examine if RASA1 is required for VV specification, we administered TM to pregnant *Rasa1^fl/fl^* mice carrying *Rasa1^fl/fl^* and *Rasa1^fl/fl^ Cdh5^ert2cre^* embryos at E12.5 and analyzed embryos at E14.5 for PROX1 and VEGFR3 expression at the respective venous junctions (Figure 1 B). Loss of RASA1 in EC at E12.5 did not affect VV specification. Therefore, RASA1 is dispensable for the specification of VV as well as LVV.

### RASA1 is required for the continued development of LVV and central VV

Development of LVV and central VV continues through E16.5 as valve leaflets are extended into the vessel lumen.^22, 23^ To determine if RASA1 is required for continued LVV and central VV development, we administered TM to *Rasa1^fl/fl^* and *Rasa1^fl/fl^ Cdh5^ert2cre^* embryos at E12.5 and analyzed embryos at E16.5. LVV and VV structure was analyzed by immunostaining of tissue sections and by SEM (Figure 2 A and B). In *Rasa1^fl/fl^* control embryos, one pair of normal LVV was identified at the vast majority of examined LV-BV junctions (Figure 2 C). In contrast, in *Rasa1^fl/fl^ Cdh5^ert2cre^* embryos, most LV-BV junctions contained no LVV and a minority contained only one LVV. None of the examined LV-BV junctions contained two LVV (Figure 2 C). Similarly, in *Rasa1^fl/fl^* control embryos, normal VV were found at all examined junctions of the IJV, EJV and SCV with the SVC (Figure 2 D and E). In contrast, in *Rasa1^fl/fl^ Cdh5^ert2cre^* embryos, development of central VV was severely impaired. In the majority of cases, all three VV were absent and in the remaining cases, only a rudimentary EJV VV could be identified whereas the IJV and SCV VV were also absent (Figure 2 D and E). Thus, although RASA1 is not required for initial specification of LVV and central VV, it is required for their continued development.

**Figure 2.**
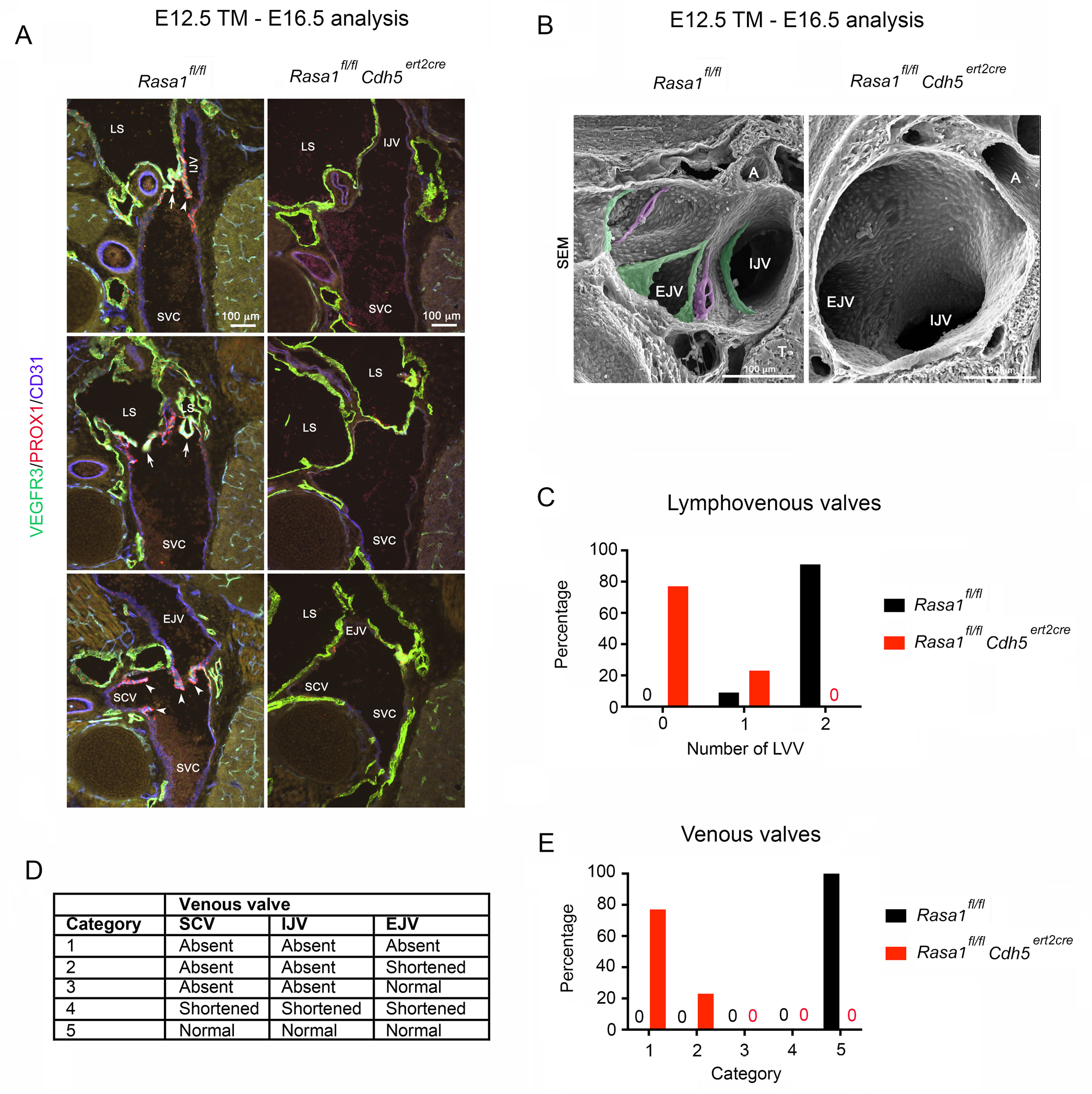
RASA1 is necessary for continued LVV and central VV development after initial specification. **A** and **B**, *Rasa1^fl/fl^* and *Rasa1^fl/fl^ Cdh5^ert2cre^* embryos were administered TM at E12.5 and harvested at E16.5. **A**, Longitudinal sections through the neck region (dorsal to ventral from top to bottom) were stained with the indicated antibodies. LS, lymphatic sac; IJV, internal jugular vein; EJV, external jugular vein; SCV, subclavian vein; SVC, superior vena cava. LVV and VV are indicated by arrows and arrowheads respectively. **B**, The structure of LVV and central VV was analyzed by SEM. A, Artery. LVV and VV are pseudo-colored in magenta and green respectively. In **A** and **B**, note absence of LVV and VV in *Rasa1^fl/fl^ Cdh5^ert2cre^* embryos. **C**, Percentage of LV-BV junctions from *Rasa1^fl/fl^* and *Rasa1^fl/fl^ Cdh5^ert2cre^* embryos in **A** and **B** that contain the indicated number of LVV. *Rasa1^fl/fl^* n=11, *Rasa1^fl/fl^ Cdh5^ert2cre^* n=13. **D**, VV structure categories. **E**, Percentage of BV junctions for *Rasa1^fl/fl^* and *Rasa1^fl/fl^ Cdh5^ert2cre^* embryos in **A** and **B** that fall into the indicated VV categories in **D**. *Rasa1^fl/fl^* n=11, *Rasa1^fl/fl^ Cdh5^ert2cre^* n=13.

### Impaired development of LVV and central VV in embryos that express RASA1 R780Q alone

Some studies have indicated that RASA1 can participate in intracellular signaling pathways independent of its ability to promote Ras hydrolysis of GTP.^6^ Therefore, we examined if the function of RASA1 in LVV and central VV development was dependent upon its catalytic GAP activity. For this purpose, we used a *Rasa1^R780Q^* allele that we reported previously.^20^ R780 of RASA1 is the “arginine finger” of the GAP domain that is essential for GAP activity.^6, 16^ Mutation of this arginine to glutamine in R780Q thus abrogates GAP activity, although is predicted to leave all other putative RASA1 functions intact. Since *Rasa1^R780Q/R780Q^* embryos die at E10.5 of gestation^16^, we generated *Rasa1^fl/R780Q^* embryos with ubiquitin promoter-driven *ert2cre* (*Ub^ert2cre^*) to examine a requirement of RASA1 GAP activity for LVV and central VV development. TM was administered to *Rasa1^fl/R780Q^ Ub^ert2cre^* embryos and control *Rasa1^fl/fl^* and *Rasa1^fl/R780Q^* embryos at E12.5 and LVV and VV development was analyzed at E16.5 as before (Figure 3 A). Compared to controls, *Rasa1^fl/R780Q^ Ub^ert2cre^* embryos showed impaired development of LVV and VV. The majority of LV-BV junctions in *Rasa1^fl/R780Q^ Ub^ert2cre^* embryos contained only one LVV (Figure 3 B). Furthermore, the majority of examined vessels from *Rasa1^fl/R780Q^ Ub^ert2cre^* embryos contained no VV at the junctions of the IJV and SCV with the SVC (Figure 3 C). Nonetheless, the impact of switch to expression of RASA1 R780Q alone upon LVV and VV development was less than that observed following complete loss of RASA1 in EC (Figure 2). This finding is consistent with the notion that RASA1 regulates LVV and VV development through both GAP domain dependent and GAP domain independent mechanisms.

**Figure 3.**
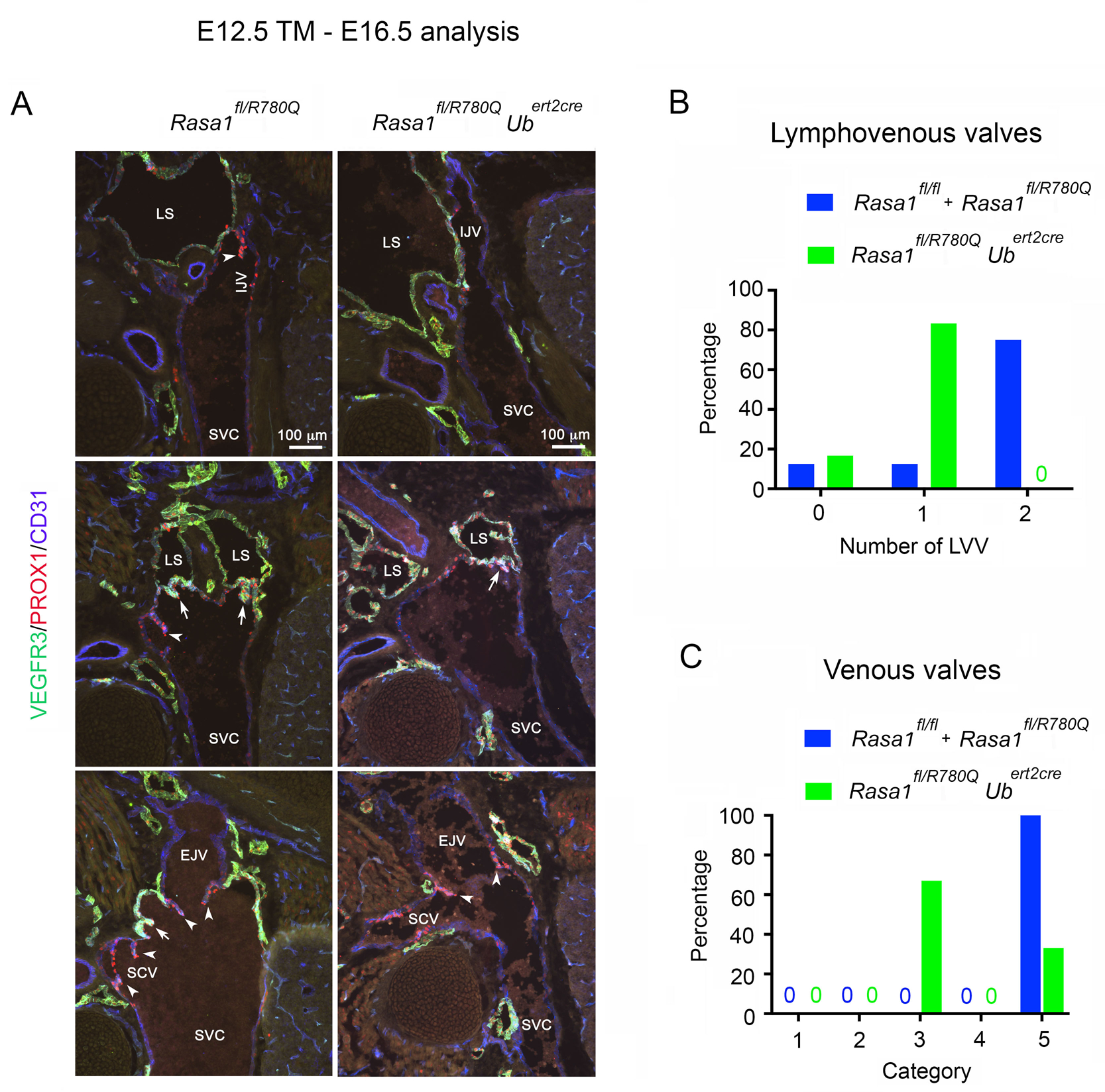
Development of LVV and central VV in RASA1 R780Q embryos. **A**, *Rasa1^fl/R780Q^* females were crossed with *Rasa1^fl/R780Q^ Ub^ert2cre^* males. TM was administered to pregnant females at E12.5 and embryos were harvested at E16.5. Longitudinal sections of the neck region (dorsal to ventral from top to bottom) were stained with the indicated antibodies. Shown are representative images from *Rasa1^fl/R780Q^* and *Rasa1^fl/R780Q^ Ub^ert2cre^* embryos. LS, lymphatic sac; IJV, internal jugular vein; EJV, external jugular vein; SCV, subclavian vein; SVC, superior vena cava. Arrows and arrowheads indicate LVV and VV respectively. **B** and **C**, Percentage of LV-BV junctions from *Rasa1^fl/R780Q^ Ub^ert2cre^* embryos and control *Rasa1^fl/fl^* and *Rasa1^fl/R780Q^* embryos from **A** that contain the indicated number of LVV (**B**) and categories of VV (**C,** see Figure 2 D). *Rasa1^fl/R780Q^ Ub^ert2cre^* n=6, *Rasa1^fl/fl^* plus *Rasa1^fl/R780Q^* n=8.

### MAPK inhibition partially rescues LVV development in the absence of RASA1

EC in developing LVV in *Rasa1^fl/fl^ Cdh5^ert2cre^* embryos administered TM at E12.5 showed increased activation of MAPK at E14.5 (Figure 4 A and B). Therefore, we next asked if inhibition of MAPK could rescue development of LVV and central VV in TM-treated *Rasa1^fl/fl^ Cdh5^ert2cre^* embryos. Thus, we administered the MAPK inhibitor, AZD6244, along with TM to *Rasa1^fl/fl^ Cdh5^ert2cre^* and *Rasa1^fl/fl^* control embryos at E12.5 and on each subsequent day thereafter until embryo harvest at E16.5. AZD6244 had a significant rescue effect upon LVV development in TM-treated *Rasa1^fl/fl^ Cdh5^ert2cre^* embryos (Figure 4 C and D). Whereas in the absence of AZD6244 treatment, the majority of LV-BV junctions contained no LVV, in the presence of AZD6244 the majority of LV-BV junctions contained at least one LVV and in some cases two LVV were identified. AZD6244 had no influence upon LVV development in TM-treated *Rasa1^fl/fl^* embryos. These findings indicate that impaired LVV development results in part from dysregulated activation of the MAPK pathway. Surprisingly, despite the finding of significant impairment of central VV development in embryos that express RASA1 R780Q alone, AZD6244 was largely unable to rescue central VV development in TM-treated *Rasa1^fl/fl^ Cdh5^ert2cre^* embryos (Figure 4 C and E). Therefore, impaired development of central VV in the absence of RASA1 may be more dependent upon dysregulated activation of distinct effector pathways downstream of activated Ras.

**Figure 4.**
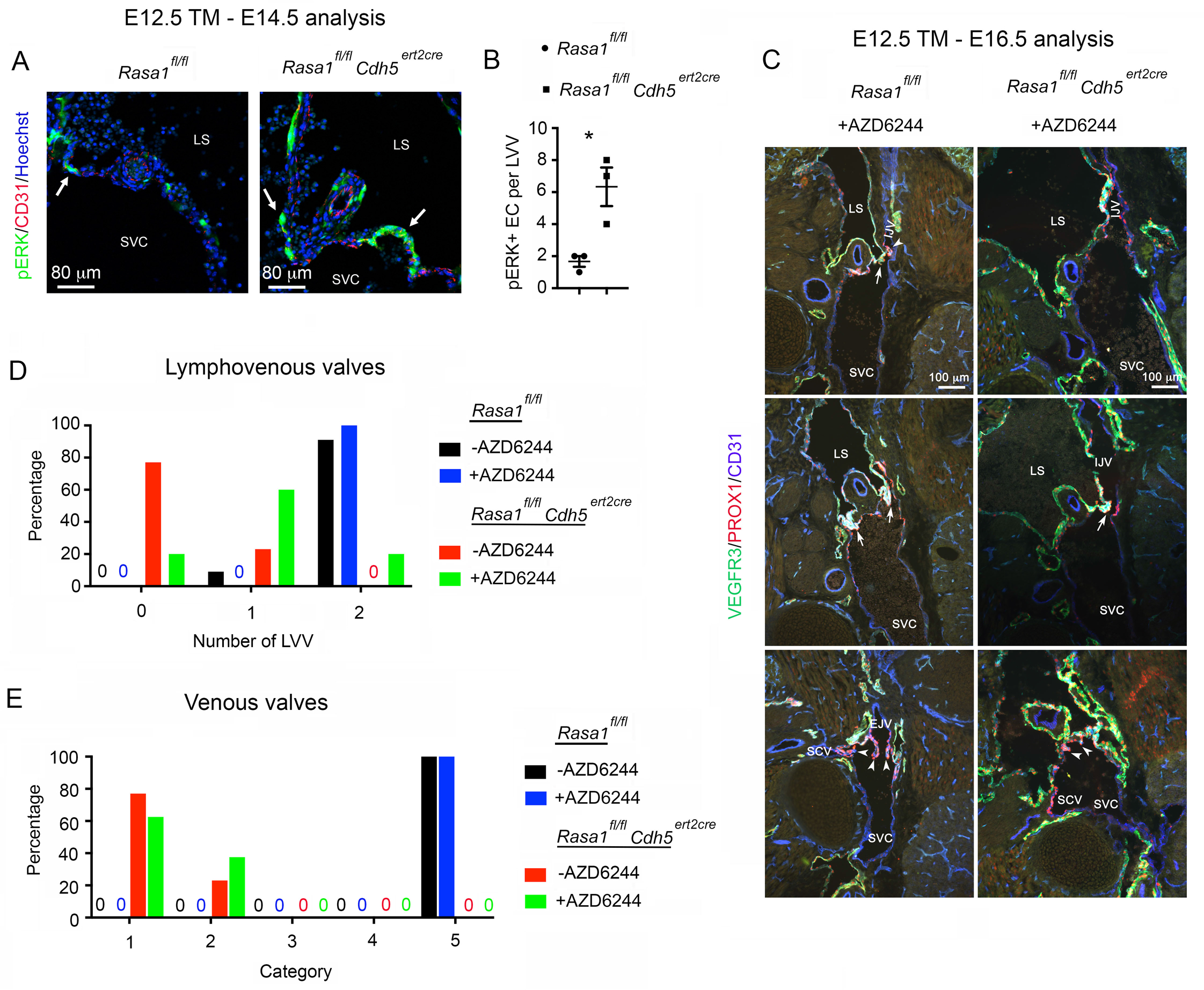
MAPK activation in LVV and effect of MAPK inhibition upon LVV and VV development in induced EC-specific RASA1-deficient embryos. **A**, *Rasa1^fl/fl^* and *Rasa1^fl/fl^ Cdh5^ert2cre^* embryos were administered TM at E12.5 and embryos were harvested at E14.5. Longitudinal sections through the neck region were stained with the indicated antibodies. Representative sections show an increased number of EC in LVV of *Rasa1^fl/fl^ Cdh5^ert2cre^* embryos with activated phospho-ERK MAPK (pERK) (arrows). LS, lymphatic sac; SVC, superior vena cava. **B**, Mean + 1 SEM of the number of pERK+ EC in LVV of embryos of the indicated genotypes (n=3). *, *P*<0.05, Student’s 2-sample t-test. **C**, *Rasa1^fl/fl^* and *Rasa1^fl/fl^ Cdh5^ert2cre^* embryos were administered TM and AZD6244 at E12.5 and AZD6244 every day thereafter until embryo harvest at E16.5. Longitudinal sections through the neck region (dorsal to ventral from top to bottom) were stained with the indicated antibodies. **D** and **E**, Percentage of LV-BV junctions from *Rasa1^fl/fl^* and *Rasa1^fl/fl^ Cdh5^ert2cre^* embryos from **C** that contain the indicated number of LVV (**D**) and categories of VV (**E,** see Figure 2 D). *Rasa1^fl/fl^* n=4, *Rasa1^fl/fl^ Cdh5^ert2cre^* n=12. Note that data from *Rasa1^fl/fl^* and *Rasa1^fl/fl^ Cdh5^ert2cre^* embryos treated with TM alone from Figure 2 C and E (red and black bars) is also plotted for comparison.

### Loss of RASA1 results in accumulation of collagen IV in EC of LVV

We recently determined that global or EC-specific disruption of *Rasa1* at E13.5 results in the apoptotic death of all dermal EC at E18.5 or E19.5 respectively.^19^ In the absence of RASA1, the vascular BM protein, collagen IV, is retained within the endoplasmic reticulum (ER) of EC and smooth muscle cells. As a result of the reduced amounts of collagen IV in BM, EC fail to attach properly to BM and undergo apoptotic death. Based upon these findings, we asked if induced loss of RASA1 at E12.5 results in intracellular accumulation of collagen IV in EC of LVV. An inability of LVV EC to export collagen IV for deposition in the ECM core of developing LVV leaflets would provide an explanation for failed LVV development. In embryos administered TM at E12.5, developing LVV are present at E14.5 (Figure 1 B) but absent at E16.5 (Figure 2). Therefore, we examined E15.5 embryos for any evidence of EC collagen IV accumulation in LVV EC. At E15.5, LVV in TM-treated (at E12.5) *Rasa1^fl/fl^ Cdh5^ert2cre^* embryos were frequently abnormal and separation of the two EC layers was often observed (Supplemental Figure 1). We examined collagen IV accumulation in embryos that additionally carried a *Prox1-eGFP* transgene. ^30^ In LVV EC of *Rasa1^fl/fl^* embryos, eGFP should be present throughout the cell cytoplasm and nucleus resulting in the appearance of a broadly distributed cellular green fluorescence signal. In contrast, in LVV EC of *Rasa1^fl/fl^ Cdh5^ert2cre^* embryos, green fluorescence signals should be visibly absent in regions of the cell cytoplasm that contain large intracellular accumulations of collagen IV, i.e. in the ER.As shown in Figure 5 and Supplemental Figure 2, in LVV of *Rasa1^fl/fl^ Cdh5^ert2cre^* embryos, individual LVV EC could be identified that contained large accumulations of collagen IV that displaced the cytoplasmic eGFP signal. No such accumulations of collagen IV were identified in LVV EC of *Rasa1^fl/fl^* embryos (Figure 5).

**Figure 5.**
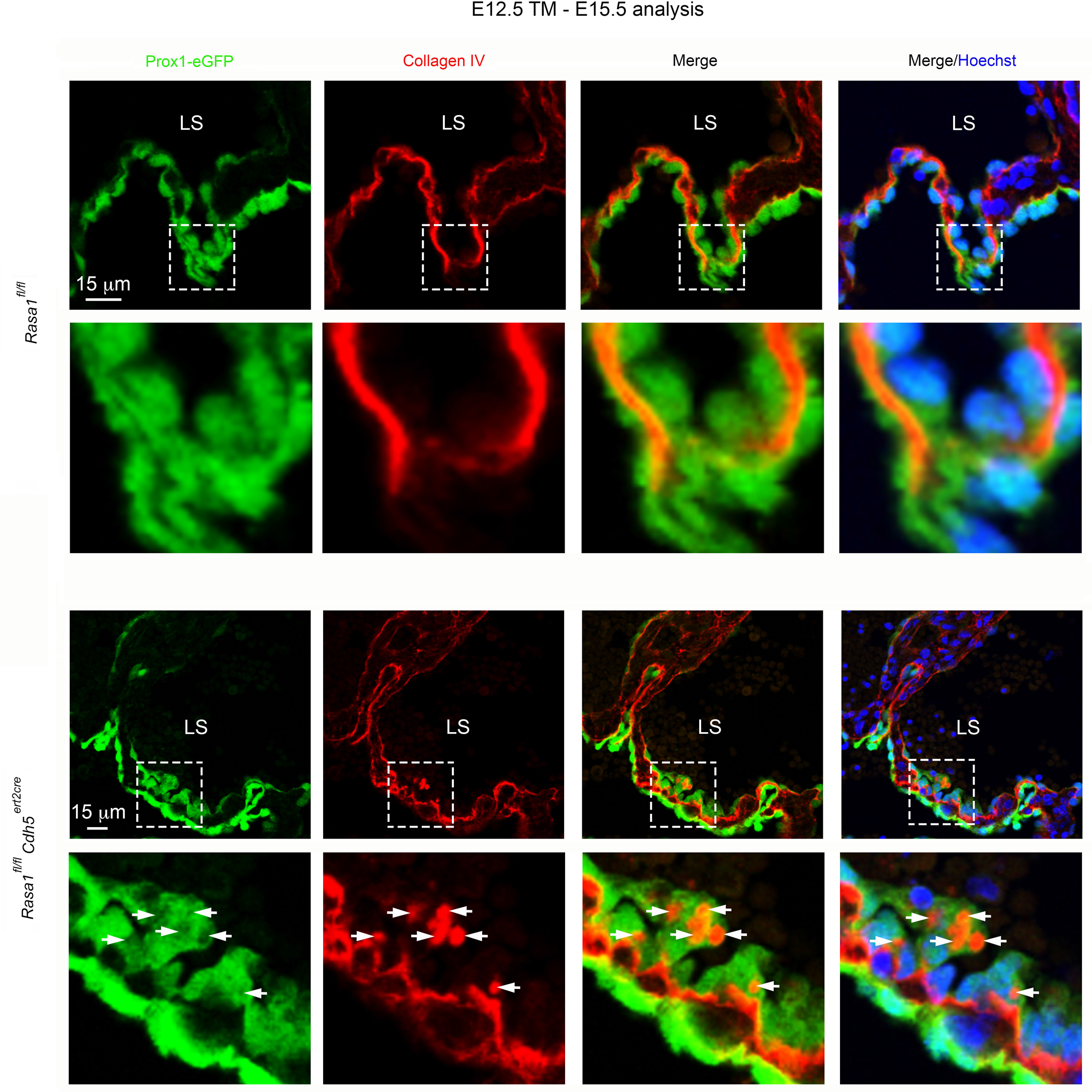
Accumulation of collagen IV in EC of developing RASA1-deficient LVV. *Rasa1^fl/fl^ Prox1-eGFP* and *Rasa1^fl/fl^ Cdh5^ert2cre^ Prox1-eGFP* embryos were administered TM at E12.5 and harvested at E15.5. Longitudinal sections through the neck region were stained with antibodies against collagen IV and the Hoechst nuclear stain. Lower power images of LVV are shown at top. Higher power images of boxed areas are shown below. LS, lymphatic sac. Note intracellular accumulation of collagen IV that corresponds to regions of displacement of the eGFP signal in EC of *Rasa1^fl/fl^ Cdh5^ert2cre^* LVV (arrows). Additional images of collagen IV accumulation in EC of LVV of *Rasa1^fl/fl^ Cdh5^ert2cre^* embryos are provided in Supplemental Figure 2.

### Rescue of LVV development by a chemical chaperone that promotes folding of collagen IV

Collagen IV becomes trapped in the ER of RASA1-deficient EC as a consequence of impaired collagen IV folding. Accordingly, 4PBA, a chemical chaperone that promotes folding of collagen IV in the ER,^31, 32^ rescues EC export of collagen IV and EC apoptosis. ^19^ Therefore, we examined if 4PBA could rescue LVV and central VV development in the absence of RASA1. 4PBA was administered along with TM to *Rasa1^fl/fl^* and *Rasa1^fl/fl^ Cdh5^ert2cre^* embryos at E12.5 and on all subsequent days until embryo harvest at E16.5. 4PBA rescued LVV development in these experiments (Figure 6 A and B). In TM plus 4PBA-treated *Rasa1^fl/fl^ Cdh5^ert2cre^* embryos, approximately half of examined LV-BV junctions contained one LVV and the remaining half contained two LVV (compared to no LVV in the majority of LV-BV junctions in *Rasa1^fl/fl^ Cdh5^ert2cre^* embryos that received TM alone). No influence of 4PBA upon LVV development was noted in TM-treated *Rasa1^fl/fl^* embryos (Figure 6 A and B). In addition, 4PBA partially rescued development of the EJV VV although was unable to rescue development of the IJV and SCV VV (Figure 6 A and C). These findings provide support for the idea that impaired development of LVV and, in part, the EJV VV is a consequence of an inability of EC to export collagen IV for deposition in the ECM core of the respective valve leaflets. Consistent with this, close inspection of 4PBA-rescued LVV revealed normal export of collagen IV by LVV EC and deposition in the LVV ECM core (Figure 6 D).

**Figure 6.**
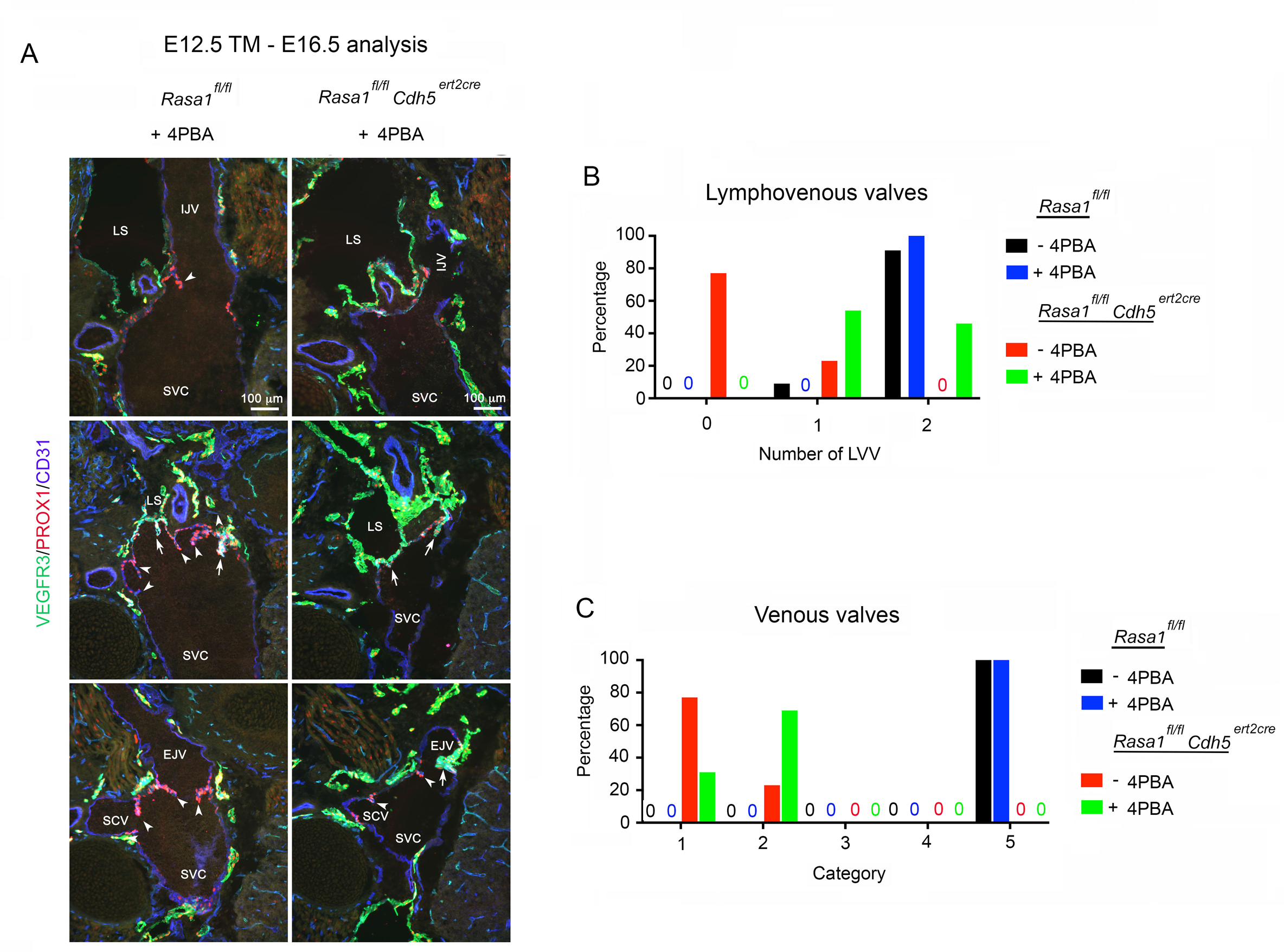
Rescue of LVV Development in induced EC-specific RASA1-deficient embryos by the chemical chaperone 4PBA. **A-D**, *Rasa1^fl/fl^* and *Rasa1^fl/fl^ Cdh5^ert2cre^* embryos were administered TM and 4PBA at E12.5 and 4PBA every day thereafter until embryo harvest at E16.5. Longitudinal sections of the neck region (dorsal to ventral from top to bottom) were stained with the indicated antibodies. LS, lymphatic sac; IJV, internal jugular vein; EJV, external jugular vein; SCV, subclavian vein; SVC, superior vena cava. In (A) arrows and arrowheads indicate LVV and VV respectively. **B** and **C**, Percentage of LV-BV junctions from *Rasa1^fl/fl^* and *Rasa1^fl/fl^ Cdh5^ert2cre^* embryos from (**A**) that contain the indicated number of LVV (**B**) and categories of VV (**C,** see Figure 2 D). *Rasa1^fl/fl^* n=4, *Rasa1^fl/fl^ Cdh5^ert2cre^* n=13. Note that data from *Rasa1^fl/fl^* and *Rasa1^fl/fl^ Cdh5^ert2cre^* embryos treated with TM alone from Figure 4 C and E (red and black bars) is also plotted for comparison. **D**, Normal deposition of collagen IV in LVV of *Rasa1^fl/fl^ Cdh5^ert2cre^* embryos treated with 4PBA.

### Rescue of LVV development by an inhibitor of collagen IV modifying enzymes

Folding of two alpha 1 and one alpha 2 collagen IV monomers into a triple helical protomer conformation in the ER is a highly regulated process that requires multiple post-translational modifications of monomers mediated by protein disulphide isomerases (PDIs), protein peptidyl isomerases (PPIs), proline-3 and proline-4 hydroxylases (P3H1-3 and P4HA1-3 respectively) and lysine hydroxylases and glycosylases (procollagen-lysine, 2-oxoglutarate 5-dioxygenases – PLODs 1-3). ^33–36^ Under- or over-post-translational modification of collagen IV monomers can affect folding of monomers into the triple helical conformation. Proteomic analyses of induced RASA1-deficient embryonic BEC established that of the P3H, P4HA and PLOD enzymes that could be detected in these cells (7 out of a total of 9), all were increased in abundance compared to control RASA1-sufficient BEC. ^19^ P3Hs, P4HAs and PLODs all belong to the same family of enzymes known as 2-oxoglutarate (2OG)-dependent oxygenases (2OG-DO) and, consequently, can be collectively inhibited by drugs such as 2,4 pyridinedicarboxylic acid (2,4 PDCA). ^37, 38^ Consistent with the notion that increased abundance of 2OG-DO in embryonic RASA1-deficient BEC is responsible for the impaired folding and export of collagen IV, 2, 4 PDCA rescued collagen IV retention and BEC apoptosis in induced global RASA1-deficient embryos. ^19^ Based on these findings, we asked if 2,4 PDCA could also rescue LVV and VV development in the absence of RASA1. As in previous drug rescue experiments, 2,4 PDCA was administered with TM to E12.5 *Rasa1^fl/fl^* and *Rasa1^fl/fl^ Cdh5^ert2cre^* embryos and on successive days thereafter until embryo harvest at E16.5 (Figure 7). 2,4 PDCA was able to rescue the impaired development of LVV in the absence of RASA1. Following drug treatment, the majority of LV-BV junctions contained 2 LVV (Figure 7 B). Furthermore, 2,4 PDCA restored export of collagen IV by LVV EC (Figure 7 D). In addition, 2,4 PDCA almost completely rescued VV development. Following drug treatment, the majority of examined BV junctions contained normal IJV, EJV and SCV VV (Figure 7 C).

**Figure 7.**
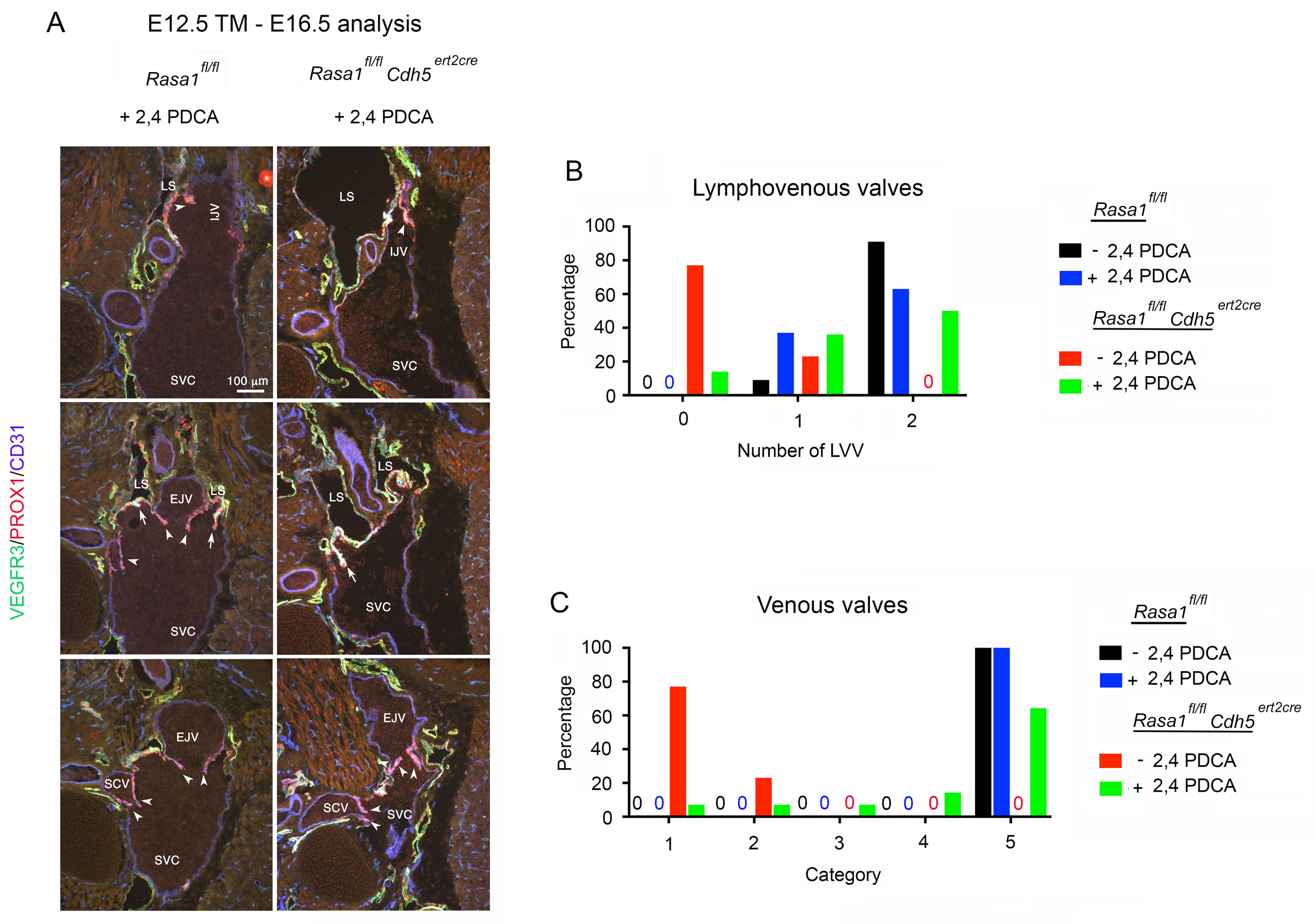
Rescue of LVV development in induced EC-specific RASA1-deficient embryos by 2,4 PDCA. **A-D**, *Rasa1^fl/fl^* and *Rasa1^fl/fl^ Cdh5^ert2cre^* embryos were administered TM and 2,4 PDCA at E12.5 and 2,4 PDCA every day thereafter until embryo harvest at E16.5. Longitudinal sections of the neck region (dorsal to ventral from top to bottom) were stained with the indicated antibodies. LS, lymphatic sac; IJV, internal jugular vein; EJV, external jugular vein; SCV, subclavian vein; SVC, superior vena cava. In (A) arrows and arrowheads indicate LVV and VV respectively. **B** and **C**, Percentage of LV-BV junctions from *Rasa1^fl/fl^* and *Rasa1^fl/fl^ Cdh5^ert2cre^* embryos from (**A**) that contain the indicated number of LVV (**B**) and categories of VV (**C,** see Figure 2 D). *Rasa1^fl/fl^* n=11, *Rasa1^fl/fl^ Cdh5^ert2cre^* n=14. Note that data from *Rasa1^fl/fl^* and *Rasa1^fl/fl^ Cdh5^ert2cre^* embryos treated with TM alone from Figure 2 C and E (red and black bars) is also plotted for comparison. D, Normal deposition of collagen IV in LVV of *Rasa1^fl/fl^ Cdh5^ert2cre^* embryos treated with 2,4 PDCA.

### RASA1 is necessary for the maintenance of central venous valves in adults

To determine if RASA1 is necessary for the maintenance of central VV in adults, we administered TM to adult littermate *Rasa1^fl/fl^* and *Rasa1^fl/fl^ Ub^ert2cre^* mice and examined VV function 3 months later. As a first approach, we examined central venous pressure that would be expected to be elevated in mice as a consequence of impaired VV function.^39^ Central venous pressure was monitored by telemetry throughout the 24-hour daily cycle for 6 days for each examined mouse (Figure 8 A). In these experiments *Rasa1^fl/fl^ Ub^ert2cre^* mice showed increased central venous pressure compared to *Rasa1^fl/fl^* controls consistent with impaired VV function.

**Figure 8.**
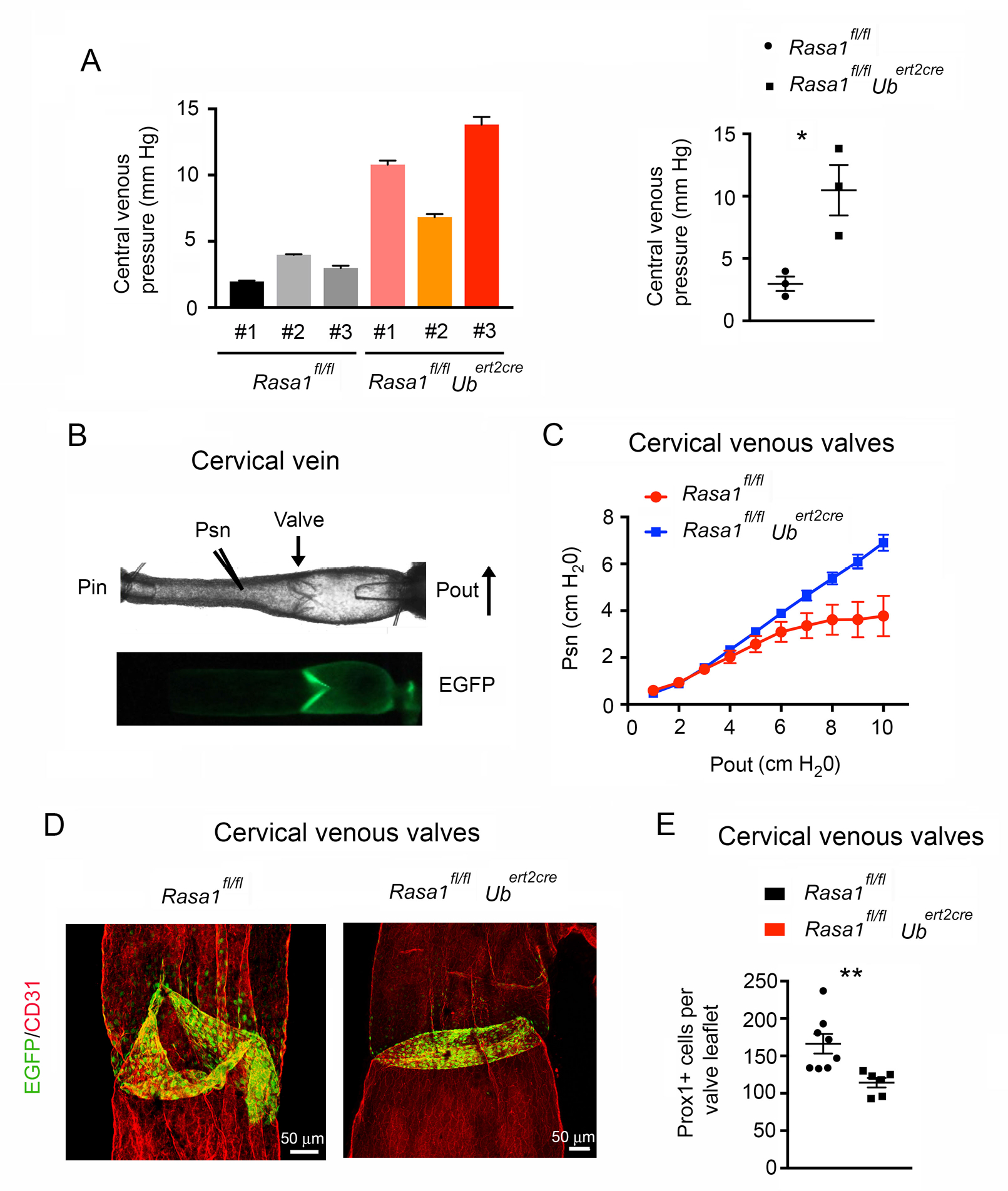
A required role for RASA1 in the maintenance of central VV function in adult mice. TM was administered to adult *Rasa1^fl/fl^* and *Rasa1^fl/fl^ Ub^ert2cre^* mice and central venous blood pressure was measured by telemetry 3 months later. At left is shown the mean + 1 SEM of the central venous pressure for 3 individual mice of each genotype based upon continuous monitoring over 6 days (see Methods). At right is shown the mean + 1 SEM of the mean central venous pressure of individual mice at left. *, *P*<0.05; Student’s 2-sample t-test. **B**, Image of a dissected cervical vein from a *Prox1-eGFP* mouse showing positions of P_in_, P_out_ and P_sn_ micropipettes in low pressure back-leak assays (top) and the fluorescent eGFP positive VV (bottom). **C**, Low pressure back-leak tests of cervical VV from adult *Rasa1^fl/fl^ Prox1-eGFP* and *Rasa1^fl/fl^ Ub^ert2cre^ Prox1-eGFP* mice administered TM 3 months previously. *Rasa1^fl/fl^ Prox1-eGFP* n=9, *Rasa1^fl/fl^ Ub^ert2cre^ Prox1-eGFP* n=8. Note leakage of *Rasa1^fl/fl^ Ub^ert2cre^ Prox1-eGFP* valves throughout the P_out_ pressure range. **D**, Whole mount images of representative cervical veins from *Rasa1^fl/fl^ Prox1-eGFP* and *Rasa1^fl/fl^ Ub^ert2cre^ Prox1-eGFP* mice in **C.** Vessels were stained with an anti-CD31 antibody prior to mounting. **E**, Mean + 1 SEM of the number of EC per VV leaflet. *Rasa1^fl/fl^ Prox1-eGFP* n=8, *Rasa1^fl/fl^ Ub^ert2cre^ Prox1-eGFP* n=6. **, *P*<0.01, Student’s 2-sample t-test.

To examine central VV function directly, cervical veins were dissected from mice and VV function was tested in low pressure back-leak assays. In these assays, veins were trimmed to contain a single VV and were cannulated at both ends to allow manipulation of intraluminal pressure upstream (P_in_) and downstream (P_out_) of the valve. For back-leak assays, P_in_ was held constant at 0.5 cm H_2_O and P_out_ was elevated over a physiological range from 0.5 to 10 cm of H_2_O. The ability of valves to resist backflow in response to elevated pressure downstream of the valve was measured with the use of a servo null micropipette (P_sn_) inserted through the vessel wall into the upstream lumen (Figure 8 B). Cervical VV from *Rasa1^fl/fl^* mice were able to resist transfer of elevated downstream pressure to the upstream vessel lumen starting at around 4 cm of H_2_O (Figure 8 C). In contrast, cervical VV from *Rasa1^fl/fl^ Ub^ert2cre^* mice were unable to prevent backflow throughout the entire 0.5 to 10 cm H_2_0 range (Figure 8 C). Thus, cervical VV are functionally impaired in *Rasa1^fl/fl^ Ub^ert2cre^* mice.

Impaired LV valve function in adult induced RASA1-deficient mice is explained by reduced cellularity of LV valve leaflets resulting in shortening of leaflets and failed LV valve closure. ^20^ Therefore, we examined the cellularity of cervical VV leaflets in induced RASA1-deficient and control mice. As determined with the use of the *Prox1- eGFP* transgene, which allows ready identification and enumeration of VV EC, induced loss of RASA1 in adult mice resulted in significantly fewer EC in cervical VV leaflets compared to controls (Figure 8 D and E). Therefore, RASA1 is necessary to maintain numbers of EC in VV leaflets which accounts for impaired VV function in induced RASA1-deficient mice

## Discussion

In this study we show that RASA1 performs a non-redundant role in the development of LVV and the development and maintenance of VV. These findings extend our knowledge of the function of this RasGAP in both blood and lymphatic vascular systems. A role for RASA1 in the continued development of LVV and VV but not their initial specification parallels our earlier reported finding regarding the role of RASA1 in LV valve development.^20^ LVV and VV development was also impaired in embryos in which RASA1 R780Q alone was expressed in EC. However, the impairment in LVV and VV development in these embryos was less than in embryos that lacked RASA1 in EC. This suggests that RASA1 regulates LVV and VV development through mechanisms that are independent of its GAP activity as well as mechanisms that are dependent upon its GAP activity. For LVV development, this is consistent with the finding that although a MAPK inhibitor was able to rescue valve development, rescue was incomplete. Moreover, for VV development, the same MAPK inhibitor was largely unable to rescue valve development in the absence of RASA1. Previously, we found that the RASA1 R780Q form of RASA1 was able to partially rescue the impaired function of adult LV valves resulting from induced loss of RASA1. ^20^ Thus, the time to development of LV valve dysfunction in adult mice induced to express RASA1 R780Q alone is substantially delayed compared to adult mice induced lose RASA1 completely. Which Ras-independent signaling pathways contribute to LVV and VV development and maintenance of mature LV valves remains to be determined.

Abnormalities of LVV are first apparent at E15.5 following disruption of *Rasa1* at E12.5. At this time, LEC and BEC layers of forming LVV detach from one another. In addition, intracellular accumulation of collagen IV can be detected in EC of valves at this time point. Intracellular accumulation of collagen IV is ultimately observed in all EC in which *Rasa1* is disrupted prior to E15.5 and results in their apoptotic death, either as a result of anoikis (detachment from the substratum) or induction of an unfolded protein response. ^19^ We propose that impaired export of collagen IV from EC of induced RASA1-deficient LVV, VV and LV valves is responsible for failed valve development. An inability of RASA1-deficient valve forming EC to export collagen IV for deposition in the developing ECM core of valves would be expected to result in EC detachment from the valve and/or induction of a UPR. Blocked export of collagen IV from RASA1-deficient embryonic EC and hemorrhage can be rescued by 4PBA that promotes folding of collagen IV in the ER. Similarly, in the current study, 4PBA largely rescued the impasse in LVV development and partially rescued the block in VV development in the absence of RASA1. These findings support the concept that collagen IV is retained within EC of developing RASA1-deficient LVV and VV because it is improperly folded. Our earlier proteomic studies of RASA1-deficient BEC identified increases in the abundance of all detectable collagen IV-modifying 2OG-DO enzymes in these cells. Evidence that this increase is responsible for collagen IV misfolding and retention in BEC was provided by the finding that drugs such as ethyl-3,4-dihydroxybenzoic acid (EDHB) and 2,4 PDCA, which broadly inhibit this class of enzymes, rescued collagen IV export from BEC and prevented hemorrhage. Similarly, as shown here, 2,4 PDCA rescued both LVV and VV development in the absence of RASA1. This is consistent with a model in which an increased abundance of 2OG-DO in EC of LVV and VV results in collagen IV retention in the ER, that in turn is responsible for blocked development of these valves in the absence of RASA1.

Induced loss of RASA1 in adult mice has not previously been reported to result in any spontaneous BV phenotypes.^18^ This can be explained on the grounds that deposition of collagen IV in BM primarily occurs during developmental angiogenesis. Collagen IV is an inherently stable molecule, thus obviating a requirement for EC to continue to engage in high rate collagen IV synthesis in postnatal life.^40^ A requirement of RASA1 for VV maintenance in adults represents an exception to this and parallels a requirement of RASA1 in the maintenance of LV valves in adults.^20^ Dysfunction of LV valves in induced RASA1-deficient mice can be explained by gradual loss of LEC from LV valve leaflets resulting in leaflet shortening below a threshold level necessary prevent to fluid backflow when closed leaflets overlap. Similarly, we show here that VV dysfunction is associated with reduced cellularity of VV leaflets. EC in LV valves and VV would be subject to higher shear stress forces than EC in LV and BV walls to the extent that continued high rate synthesis of collagen IV may be necessary for these cells to remain attached to the valve leaflet. ^41, 42^ Indeed, it is established that EC synthesize higher amounts of collagen IV when subject to shear stress in vitro.^43^ In this regard, induced loss of RASA1 in adult LV valve and VV EC may result in their detachment from leaflets or apoptotic death consequent to induction of a UPR, which may responsible for the leaflet shortening over time.

Last, based upon the studies reported herein, it is conceivable that in CM-AVM, acquired somatic second hit mutations of the *RASA1* gene in LVV-forming and VV-forming cells or their precursors could result in impaired development and function of affected LVV and VV in these patients. Whether or not CM-AVM patients show abnormalities of LVV and VV function is currently unknown.

## Methods

### Mice

*Rasa1^fl/fl^* and *Rasa1^fl/R780Q^* mice with and without *Ub^ert2cre^* transgenes have been described.^16–18, 20^ *Cdh5^ert2cre^* mice were obtained from Cancer Research UK.^29^ *Prox1- eGFP* mice were obtained from Dr. Young Kwon Hong at the University of Southern California. *Rasa1^fl/fl^ Cdh5^ert2cre^* and *Rasa1^fl/fl^ Prox1-eGFP* mice with and without *Cdh5^ert2cre^* and *Ub^ert2cre^* were generated through cross-breeding. All mice were on a mixed 129S6/SvEv X C57BL/6 genetic background. All experiments performed with mice were in compliance with University of Michigan and University of Missouri guidelines and were approved by the respective university committees on the use and care of animals.

### LVV and VV specification and development

Pregnant *Rasa1^fl/fl^* mice carrying *Rasa1^fl/fl^* embryos with and without *Cdh5^ert2cre^* and pregnant *Rasa1^fl/fR780Q^* mice carrying *Rasa1^fl/fl^*, *Rasa1^fl/R780Q^* and *Rasa1^R780Q/R780Q^* embryos with and without and *Ub^ert2cre^* were given 2 i.p. injections of TM as above on consecutive days starting at E10.5, E12.5 or E14.5 of gestation. For some pregnant mice, AZD6244 (Selleckchem; 0.05 mg/g body weight per injection), 4-phenylbutyric acid (4PBA; Sigma; 0.25 mg/g body weight per injection), or 2,4 PDCA (Sigma; 0.1 mg/g body weight per injection) was injected i.p. at the same time as TM and on subsequent days until embryo harvest. Embryos were harvested at different times and fixed in 4% paraformaldehyde overnight. For embryos that were harvested at E14.5 and later, an abdominal incision was made and blood was drained from embryos before fixation.

Immunohistochemistry on sections was performed as described.^22^ Fixed embryos were embedded in OCT solution (Sakura, Tokyo, Japan) and 12 μm thick cryosections were prepared in the frontal orientation using a cryotome (Thermo Fisher Scientific, Model: HM525 NX). Primary antibodies used were: goat anti-VEGFR3 (R&D Systems), rabbit anti-PROX1 (AngioBio), rat anti-CD31 (BD Pharmingen), goat anti-collagen IV (1340- 01, Southern Biotech), rabbit anti-pERK (D13.14.4E, Cell Signaling Technology). Secondary antibodies were fluorochrome-labeled species specific anti-Igs. The antibodies were diluted in blocking buffer (PBS+ 0.1% Triton X-100 +0.1% BSA) and immunohistochemistry was performed using standard protocols. After mounting, the sections were visualized with Eclipse 80i microscope (Nikon, Tokyo, Japan) equipped with Zyla sCMOS camera (Andor Technology, Belfast, UK) and analyzed using NIS-Elements BR software (Nikon, Tokyo, Japan). Several consecutive sections were analyzed to determine the presence or absence of LVVs and VVs.

SEM was performed according to our previous protocol.^22, 44^ Briefly, 500 µm sections were prepared using vibratome (Leica, Buffalo Grove, IL, USA). Sections were fixed in 2% glutaraldehyde in 0.1 M cacodylate buffer for 2 hours. After washing profusely in 0.1M cacodylate buffer, the sections were post fixed in 1% osmium tetroxide in 0.1 M cacodylate buffer for 2 hours and subsequently dehydrated in a graded ethanol series. The sections were further dehydrated in hexamethyldisilazane and allowed to air-dry overnight. Dry sections were sputter-coated with Au/Pd particles (Med-010 Sputter Coater by Balzers-Union, USA) and observed under Quanta SEM (FEI, Hillsboro, OR, USA) at an accelerating voltage of 20KV.

### Central venous pressure

Three-month old littermate *Rasa1^fl/fl^* and *Rasa1^fl/fl^ Ub^ert2cre^* mice were given 2 i.p. injections of TM (0.05 mg/g body weight per injection, dissolved in corn oil) on consecutive days. One week later, central venous pressure was determined by radiotelemetry using an implantable microminiaturized electronic monitor (PA-C10; Data Sciences International) as described with minor modifications.^45^ Briefly, the catheter of the device was passed into the right EJV and the transducer was placed in the abdominal cavity. After surgery, mice were individually housed in cages atop receiver pads allowing for real-time measurements of venous pressure, heart rate and activity. To minimize the influence of fluctuations in venous pressure resulting from changes in respiration and activity, for each mouse an average central venous pressure was calculated based upon readings taken every 10 seconds for 15 minutes every 3 hours for 6 days total.

### Vein dissection and cannulation

Three-month old littermate *Rasa1^fl/fl^ Prox1-eGFP* mice with and without *Ub^ert2cre^* were given 2 i.p. injections of TM as above. After 12 weeks, mice were anesthetized with sodium pentobarbital (60 mg/kg, i.p.) and placed on a heating pad. The skin under the chin and upper chest was shaved and a ventral incision was made to expose the underlying cervical and EJV in association with superior cervical afferent LV. The LV and attached connective tissue were removed and the EJV was cut centrally, held with fine forceps and pulled caudally while trimming along the sides of the vessel and its major branch, the cervical vein (and sometimes also other minor branches), until a segment ∼3-4 mm in length was obtained. Some vessels were unbranched (Figure 8 B) but in most cases the cervical vein joined the external jugular vein just distal to the valve. After cutting the distal ends of the main vessel and any branches, the entire segment was removed and placed in room temperature Krebs-BSA solution [146.9 mM NaCl; 4.7 mM KCl; 2 mM CaCl_2_·2H_2_O; 1.2 mM MgSO_4_; 1.2 mM NaH_2_PO_4_·H_2_O; 3 mM NaHCO_3_; 1.5 mM sodium-Hepes; 5 mM D-glucose; 0.5% BSA (pH 7.4 at 37◦C)] in a dissection dish with a 1 cm layer of Sylgard (Dow Corning). The mouse was then euthanized with an injection of KCl (0.2 M, i.c.). After pinning the segment with pieces of 40 µm wire, the remaining connective tissue and fat were removed and the segment was transferred to a 3 ml cannulation chamber. The proximal and distal ends were cannulated onto 90 µm glass micropipettes filled with Krebs-BSA and tied with 12-0 suture. After pressurization most of the red blood cells were flushed out and any side branches were identified and tied with 12-0 suture. The cannulation chamber, with attached pipette holders and vessel, were transferred to the stage of an inverted microscope, where the segment was heated to 37°C and perfused with Krebs buffer (0.5 ml/min). Inflow and outflow pressures were set initially using standing fluid reservoirs and the axial length was adjusted to remove slack with luminal pressure briefly set to 10 cmH_2_O. The segment was then equilibrated for 30 minutes at 2 cm H_2_O luminal pressure. Development of spontaneous tone confirmed vessel viability but the venous valve tests described below were performed after perfusion of the segment with Ca^2+^-free Krebs (Krebs with 3 mM EDTA replacing CaCl_2_·2H_2_O) for at least 20 minutes to eliminate spontaneous tone. The vessel image was digitized using a fire-wire camera (model A641FM; Basler) and inner diameter was continuously tracked using a custom computer algorithm.^46^ All valve tests were recorded as AVI files, with embedded pressure and diameter data, for later replay and valve tracking, as needed.

### Valve back-leak assays

To conduct tests of valve function, the pressure control for each cannulation pipette was switched from the reservoirs to a two-channel pump controller (Cardiovascular Research Institute, Texas A&M University) driven through a D-A interface by a LabVIEW program (National Instruments) running under Windows 7. A servo-nulling micropipette with tip diameter ∼3 µm was inserted through the vessel wall on the upstream side of the valve to measure the local pressure (Psn) at that site.^46^ Psn, inflow pressure (Pin), outflow pressure (Pout) and diameter were recorded at 30 Hz using a model PCI 6030e A-D interface (National Instruments). After micropuncture Pin and Pout were raised simultaneously from 0.5 to 10 cm H_2_O in order to verify the calibration of the servo-nulling system. To assess the ability of a closed valve to prevent pressure back leak, Pin was set to 0.5 cm H_2_O and Pout was elevated ramp-wise from 0.5 to 10 cm H_2_O over a 2-min period. The servo-nulling pipette detected any rise in pressure upstream from the valve. The Pout ramp was repeated 3 times. For data analysis Psn and Pout values were binned in 1 cm H_2_O intervals for each ramp, using a LabVIEW program, to obtain average values of Psn and Pout for plotting and statistical tests.

### Venous valve whole mount staining

Vessels from VV function assays were fixed in 1% paraformaldehyde overnight, blocked by incubation in PBS/10% donkey serum/0.3% Triton-X100 and incubated overnight with rat anti-CD31 in PBS/10% donkey serum/0.3% Triton-X100. Vessels were subsequently incubated with donkey anti-rat Ig coupled to Alexa Fluor 594 (Jackson Immunoresearch) in PBS for 2 hours before viewing on a Leica SP5 X confocal microscope. PROX1-positive EC in VV leaflets were enumerated with the use of Imaris software (Bitplane).

### Statistical analysis

*P* values were calculated using two-tailed Student’s 2-sample 2- sided *t*-testS.

### Data availability

The source data underlying all figures is provided as a Source Data File.

## Acknowledgements

We acknowledge the assistance of the Michigan Medicine Physiology Phenotyping Core for measurement of central venous pressure in mice. This work was supported by National Institutes of Health grants HL120888 and HL146352 to PDK, HL131652 and HL133216 to SS, HL-120867 to MJD and GM103441 to XG (PI: Dr. McEver).

## Author contributions

All authors designed experiments and analyzed data; D.C., X.G. P.E.L. and M.J.D. conducted experiments; P.D.K. wrote the manuscript with input from other authors.; P.D.K. supervised the project.

---

**Supplemental Figure 1.** Abnormalities of LVV structure following induced loss of RASA1. *Rasa1^fl/fl^* and *Rasa1^fl/fl^ Cdh5^ert2cre^* embryos were administered TM at E12.5 and harvested at E15.5. Longitudinal sections through the neck region were stained with the indicated antibodies. LS, lymphatic sac; IJV, internal jugular vein; SCV, subclavian vein; SVC, superior vena cava. Note separation of EC layers of LVV in *Rasa1^fl/fl^ Cdh5^ert2cre^* embryos (arrows).

**Supplemental Figure 2.** Accumulation of collagen IV in EC of developing RASA1- deficient LVV. *Rasa1^fl/fl^ Cdh5^ert2cre^ Prox1-eGFP* embryos were administered TM at E12.5 and harvested at E15.5. Longitudinal sections through the neck region were stained with antibodies against collagen IV and the Hoechst nuclear stain. Lower power images of LVV are shown at top. Higher power images of boxed areas are shown below. LS, lymphatic sac. Note intracellular accumulation of collagen IV that corresponds to regions of displacement of the eGFP signal in EC of LVV (arrows).

